# A general technique for the detection of switch-like bistability in chemical reaction networks governed by mass action kinetics with conservation laws

**DOI:** 10.1101/2020.11.06.372235

**Authors:** Brandon C Reyes, Irene Otero-Muras, Vladislav A Petyuk

**Affiliations:** Advanced Computing, Math, and Data Division, Pacific Northwest National Laboratory, Richland, Washington, 99352, United States; BioProcess Engineering Group, IIM-CSIC (Spanish National Research Council), 36208, Vigo, Spain; Biological Sciences Division, Pacific Northwest National Laboratory, Richland, Washington, 99352, United States

**Keywords:** Systems Biology, Signaling pathways, Chemical Reaction Network Theory, Mass action kinetics, Bistability, Switch-like behavior

## Abstract

**Background:** Theoretical analysis of signaling pathways can provide a substantial amount of insight into their function. One particular area of research considers signaling pathways capable of assuming two or more stable states given the same amount of signaling ligand. This phenomenon of bistability can give rise to switch-like behavior, a mechanism that governs cellular decision making. Investigation of whether or not a signaling pathway can confer bistability and switch-like behavior, without knowledge of specific kinetic rate constant values, is a mathematically challenging problem. Recently a technique based on optimization has been introduced, which is capable of finding example parameter values that confer switch-like behavior for a given pathway. Although this approach has made it possible to analyze moderately sized pathways, it is limited to reaction networks that presume a uniterminal structure. It is this limited structure we address by developing a general technique that applies to any mass action reaction network with conservation laws.

**Results:** In this paper we developed a generalized method for detecting switch-like bistable behavior in any mass action reaction network with conservation laws. The method involves 1) construction of a constrained optimization problem using the determinant of the Jacobian of the underlying rate equations, 2) minimization of the objective function to search for conditions resulting in a zero eigenvalue 3) computation of a confidence level that describes if the global minimum has been found and 4) evaluation of optimization values, using either numerical continuation or directly simulating the ODE system, to verify that a bistability region exists. The generalized method has been tested on three motifs known to be capable of bistability.

**Conclusions:** We have developed a variation of an optimization-based method for discovery of bistability, which is not limited to the structure of the chemical reaction network. Successful completion of the method provides an S-shaped bifurcation diagram, which indicates that the network acts as a bistable switch for the given optimization parameters.

## 1 Background

Cellular decisions via signaling pathways are essential for complex biological systems to function. The key attribute of signaling pathways which are capable of mediating a decision-making process is switch-like behavior. Such behavior assumes that the system has (typically) two stable equilibria and there is a way to switch between them. In bistable switches, two different thresholds for switching back and forth ensure the robustness of the decision. This characteristic dose-response behaviour is called hysteretic. Distortion of these signaling pathways with switch-like behavior manifests in no switching at all, switching at incorrect input signals, irreversible switching, etc. These malfunctions result in incorrect cell decisions and may be one of the underlying causes of developmental disorders, cancer, diabetes, and presumably a number of other pathologies [1, 2, 3]. Given that cellular decisions can consist of an immense number of biochemical interactions, smaller network motifs are often considered and can help elucidate the portions of the signaling pathway that are key to the decision-making process [4].

Although discovering essential network motifs can provide a wealth of information, obtaining these configurations is not only difficult, but costly if approached purely from an experimental point of view. For this reason, it is imperative for this process to be coupled with mathematical modeling, which can act as a guide for designing experiments. Analysis of these models is useful as they can determine the existence of bistability in the signaling pathway, an attribute directly tied to the pathway’s ability to exhibit switch like behavior. Existence of bistability in chemical reaction networks has been an active area of research since the 1970s. In particular, a wealth of mathematical theory has been oriented towards network motifs that utilize mass action kinetics for the participating reactions. This is due to the fact that mass action law does not employ assumptions on time-and concentration-scale separation as do other kinetics, such as Michaelis-Menten [5, 6, 7, 8].

One well-established theoretical framework to preclude multistationarity (and therefore bistability) in a network motif following mass action kinetics was developed by Feinberg, Horn, and Jackson [9, 10]. This theory, aptly named Chemical Reaction Network Theory (CRNT) uses the underlying structure of the reactions in the network to identify key properties. CRNT has produced results such as the Deficiency Zero and One Theorems which preclude bistability for certain network structures, irregardless of the kinetic constant values [9]. Although these theorems are very powerful, it is often the case that more complex networks found in cell signaling do not meet the deficiency requirement of these theorems [11]. To consider networks with higher deficiency, the Deficiency One Algorithm and Advanced Deficiency Algorithm were developed [12, 11]. These algorithms use the structure of the network to construct a system of equalities and inequalities. If these systems are solvable by either linear or nonlinear programming, they can state the existence of multiple positive steady states (note that a multistationary system is not necessarily bistable). In addition to these methods, injectivity theory and network concordance tests have also been developed, which attempt to address those networks that are not covered by the aforementioned theory [13, 14].

Besides CRNT and linear/nonlinear programming, there are a number of alternative approaches for gaining insights into an ODE system’s behavior. For example, algebraic methods utilizing the Gröbner basis approach [15, 16] and cylindrical algebraic decomposition [17] have been developed in concert with CRNT. The goal of which is to obtain analytical solutions to the system of ODEs at the steady state and then further analyze the solutions to detect bistability. Recently, the optimization-based approach of [18] has gained attention as it provides an efficient procedure to search for bistability with respect to the conservation laws of a mass action system. For brevity we will refer to this method as the uniterminal approach (where the term uniterminal is defined in the Basic notation section) as the approach is limited to uniterminal networks. This particular approach attempts to find bistability by constructing an efficient optimization problem to find a saddle-node and then uses numerical continuation to confirm if the saddle-node is a saddle-node bifurcation (in general for signaling pathways under study bistability occurs via saddle-node bifurcations). This approach is of considerable interest because it allows 1) evaluation of fairly large pathways and 2) one to directly put bounds on values of the parameters of the network, such as the species’ concentrations and reactions. The uniterminal approach is constructed based on the assumption that 1) every reaction is endowed with mass action kinetics 2) the network admits a strictly positive steady state (where the concentrations of all the species are positive) and 3) the network is uniterminal.

Although this optimization approach in combination with hybrid optimization solvers [19] works quite efficiently, it is limited to networks that are uniterminal. This constraint is a direct result of assumptions made in the formulation of the optimization problem. More specifically, by assuming that the network is uniterminal, the approach is able to form a basis for the deficiency subspace see section “Deficiency and equilibrium manifold” in [18]. A consequence of this result is the ability to form a square system of equations that define the equilibrium manifold of the ODE system that is compatible with the reaction polyhedron [20]. Furthermore, since this system is square, the approach can construct sufficient conditions for a saddle-node in the presence of mass conservation by minimizing the system’s determinant (a scalar value available for only square systems) see section “Sufficient conditions for a saddle-node in presence of mass conservation” in [18]. To address this limiting condition and extend the reach of this optimization approach, we constructed a general technique that can investigate the bistability of reaction networks regardless of their terminality.

The general approach (as depicted in Figure 1) uses the network structure to construct an optimization problem that searches for a saddle-node. This optimization problem differs from the uniterminal approach as it searches for a saddle-node using direct ODE stability analysis, rather than relying on constrictive assumptions. In essence, the optimization problem formed contains an objective function that results in a Jacobian with at least a zero eigenvalue when minimized to zero. Once a set of parameters conferring a saddle-node is found by this minimization, then production of a bifurcation diagram is attempted in one of two ways. One way is based on a well-established numerical continuation technique. Alternatively, the ODE system can be directly simulated from different initial conditions to compute the dose-response curve. In this case the dose-response curve serves as a surrogate of a bifurcation diagram except it does not contain an unstable branch. This option, although more computationally intensive, allows one to determine if the system is bistable in cases where numerical continuation is not possible due to an ill-conditioned Jacobian.

**Fig. 1:**
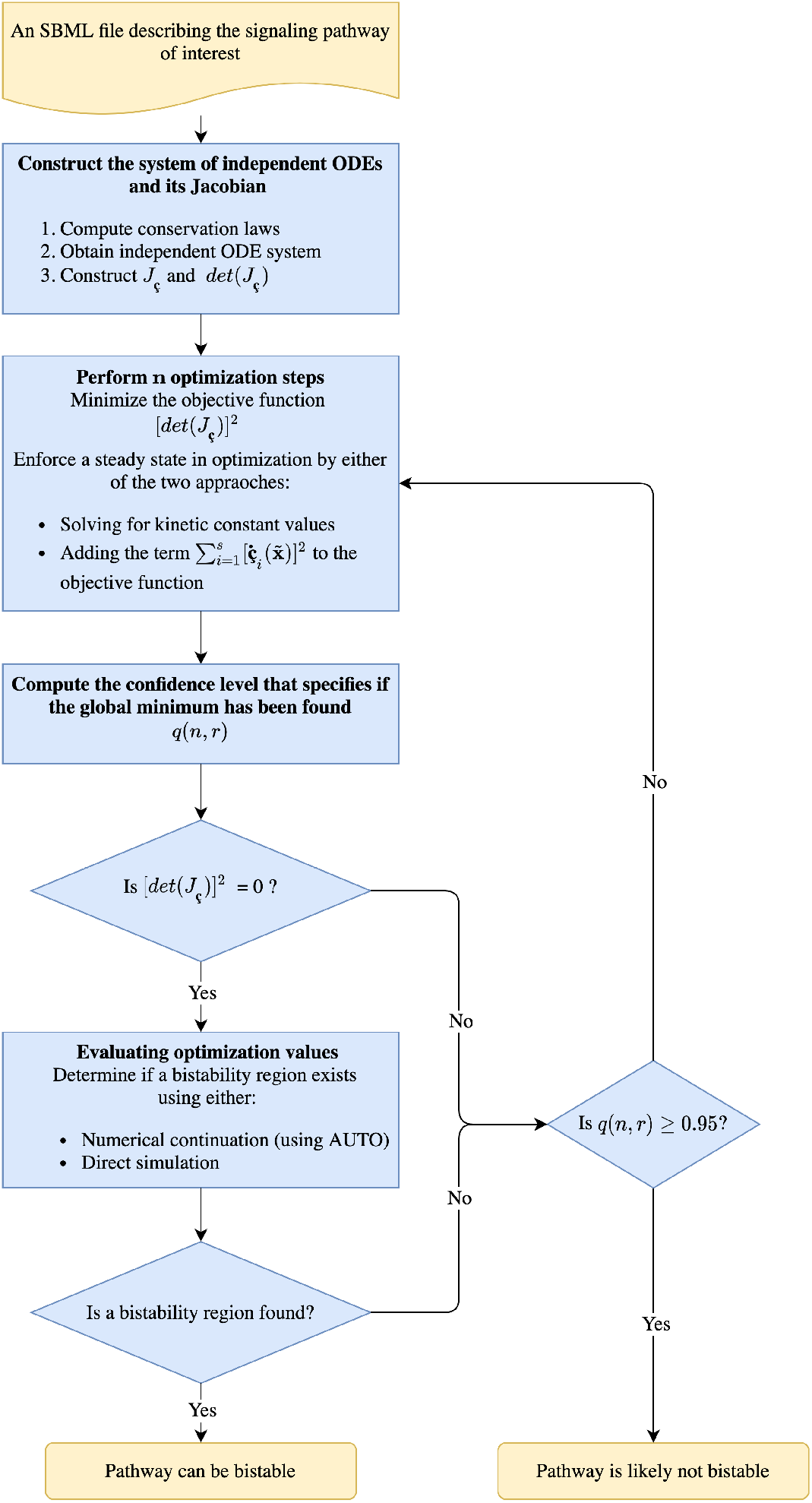
Workflow of the general approach for bistability detection in mass action chemical reaction networks. The approach is based on finding conditions that produce a Jacobian evaluated at a steady state, which has one zero eigenvalue. The symbol **ç** denotes the independent species’ concentrations. A confidence level defining if a global minimum of the optimization problem has been found is denoted as *q*(*n, r*).

### 1.1 Basic notation

Assuming mass action kinetics, a given chemical reaction network can be represented as a system of autonomous ODEs composed of *N* species and *R* reactions

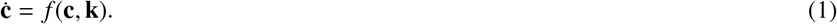

Here **ċ** denotes the temporal derivative of the species concentrations 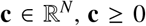, with each species having concentration *c_i_* for *i* = 1,…, *N*. The vector 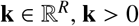, represents the kinetic rate constants of the reactions (determined by mass action kinetics), where the individual kinetic rate constants are denoted as *k_i_* for *i* = 1,…, R. We also denote the individual reactions as *r_i_* for *i* = 1,…, *R*. In addition to the system of ODEs, we also require that the network have one or more conservation laws, which we denote as follows:

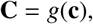

where 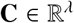, *λ* denoting the number of conservation laws, and each conservation law is given as *C_i_* for *i* = 1,…, *λ*.

A network can be described further by considering the complexes of the network. The complexes are the sets of distinct reactants or products for each reaction, and are denoted as *C_i_* for *i* = 1,…, *M*. The graph formed by the complexes and reactions of a network is then composed of disconnected subgraphs called linkage classes. For a given network we let *ℓ* be the number of linkage classes and denote them as 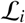 for *i* = 1,…, *ℓ*. By inspecting a linkage class further, one can then determine if a given network is uniterminal using Definitions 1, 2, and 3. The simplest depictions for a network being uniterminal and biterminal are depicted in Figure 2. Note that in the uniterminal case a linkage class containing a single complex is itself uniterminal as a single complex is strongly linked to itself [9]. Although this is assumed by CRNT, this is of little or no interest in application (as it essentially means no reaction occurs) and is simply stated for theoretical completeness.

**Fig. 2:**
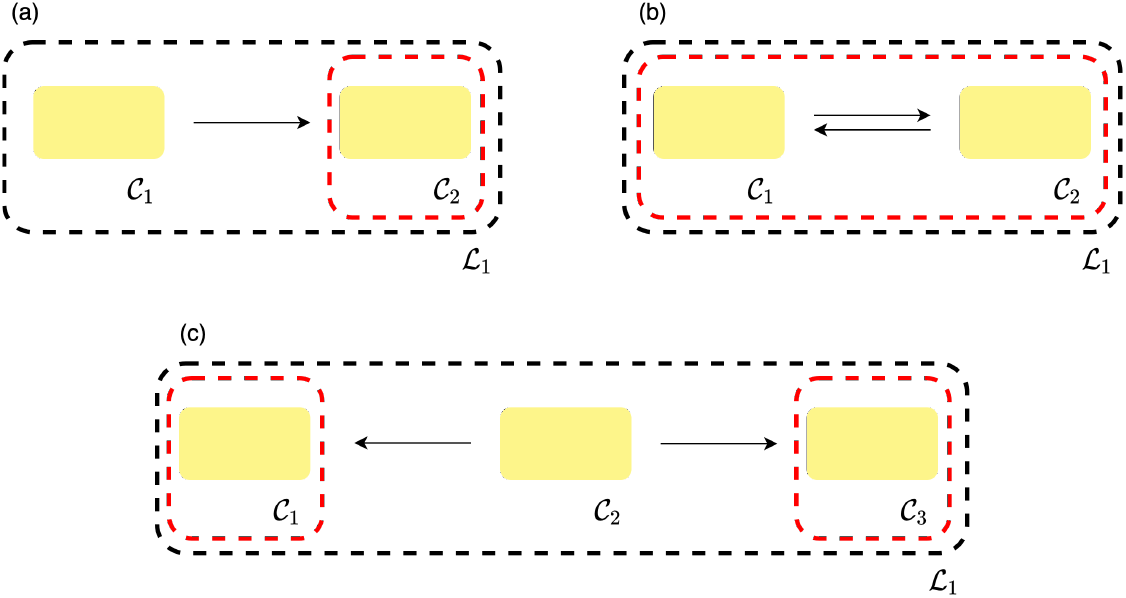
Representations of example uniterminal and biterminal networks. (a) and (b) represent the simplest possible networks that are uniterminal and (c) depicts the simplest biterminal network. The black and red dashed lines in the subfigures indicate the linkage class and terminal strong linkage classes of the network, respectively.

#### Definition 1

*Strongly linked nodes*

*Two nodes C_i_, C_j_ are said to be strongly linked if there is a directed path from C_i_ to C_j_ and also a directed path from C_j_ to C_i_*.

#### Definition 2

*Terminal strong linkage class*

*A terminal strong linkage class is a maximal set of nodes within a disconnected subgraph such that there is no edge pointing to any other set of nodes that are strongly linked*.

#### Definition 3

*Uniterminal network*

*We say that a network graph is uniterminal if every linkage class in the graph contains only one terminal strong linkage class*.

### 1.2 Bistability in Reaction Networks

Provided system (1), it’s possible that the chemical reactions will resolve to different equilibrium states depending on the starting conditions. Intuitively, such non-linear phenomenon can occur when the equilibrium concentration of one species is a polynomial function with degree greater than one with respect to an individual species. In biological systems, this behavior may present itself in a situation where a particular value of a signal species produces more than one solution with respect to a response species. In particular, bistability resulting from two stable branches connected by an unstable branch mimics switch-like action. Identification of such behavior is the focus of this manuscript.

Determining the stability of an ODE system is achieved by considering the eigenvalues of the Jacobian at a steady state point. The Jacobian of the ODE system is a matrix representation of the first-order partial derivatives of the ODE system with respect to state variables. By constructing the Jacobian one can approximate any ODE system with a system of linear ODEs at a given point. The stability at the steady state is defined by the sign of the real part of the Jacobian’s eigenvalues. If any of the eigenvalues have a positive real component, the deviation from the steady state will increase in time, which implies an unstable steady state. In the case where all of the eigenvalues have a negative real component, the deviation decreases with time, indicating a stable steady state. In some cases, such as a bistable system, a single zero eigenvalue indicates that a steady state is momentarily transitioning from stable to unstable or vice versa, which as a result of varying a parameter of the ODE system.

In a mathematical sense, this transition from a stable state to an unstable state can be exhibited by a saddle-node bifurcation. Sufficient conditions for a saddle-node bifurcation are given by Theorem 1, where the point must also be a saddle-node according to Definition 4. It should be noted that over the years the naming of a saddle-node bifurcation point has become somewhat convoluted, in other literature it is also referred to as a turning point, fold bifurcation point, or limit point bifurcation [21]. In a bistable system, two saddle-node bifurcation points delimit the range of the signal (or bifurcation parameter) for which two stable branches of steady states exist. It is this bistability phenomenon that we are particularly interested in identifying. An example scenario for bistability is depicted in Figure 3, where the parameter being varied for the bifurcation analysis is the signal of the reaction network, the solid blue line denotes a stable branch, and the dashed blue line represents an unstable branch.

**Fig. 3:**
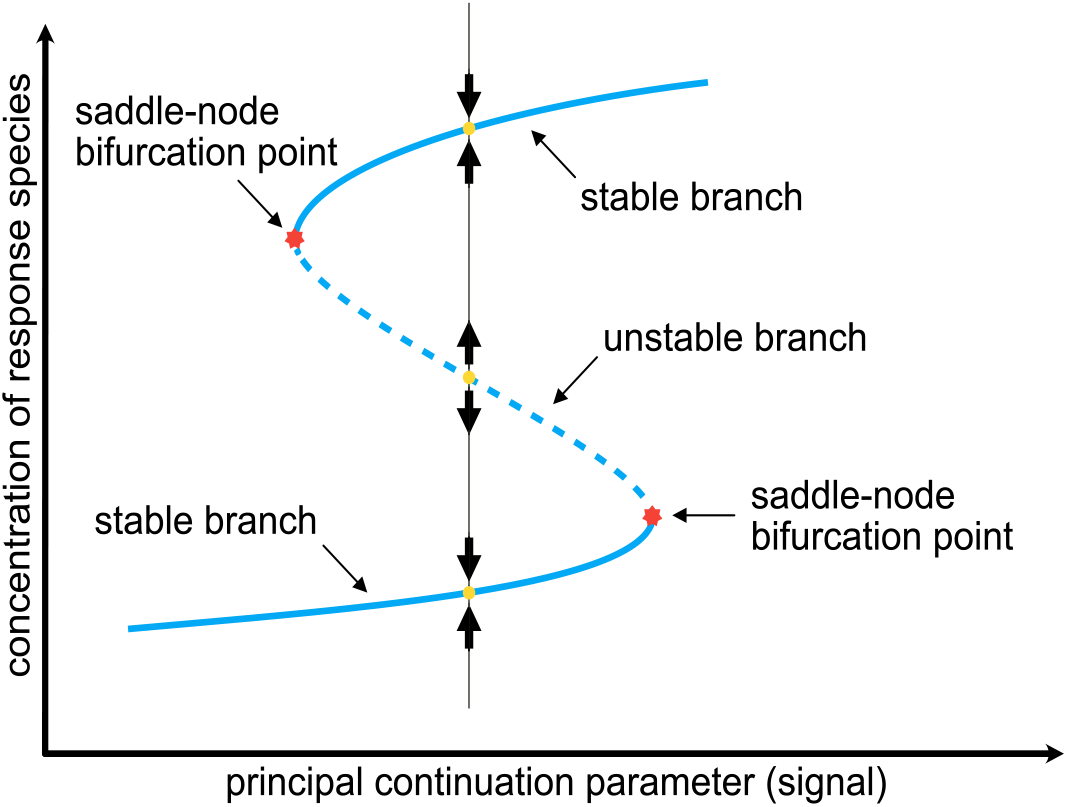
Bifurcation diagram illustrating bistability. The solid blue and dashed blue lines indicate stable and unstable branches, respectively. The figure also highlights key characteristics of bistability such as a stable and unstable point and a saddlenode bifurcation.

#### Definition 4

([22]) *Saddle-node*

*When considering an n–dimensional system of ODEs, f, we say that* 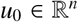 *is a saddlenode for* 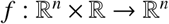 *at ϵ*_0_ *if f*(*u*_0_, *ϵ*_0_) = 0, *the linear transformation* 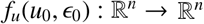 *has zero as an eigenvalue with algebraic multiplicity of one, and all other eigenvalues have nonzero real parts*.

#### Theorem 1

([22]) *Saddle-node Bifurcation Theorem*

*Suppose that* 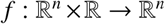 *is a smooth function, u = u*_0_ *is a saddle-node for f at ϵ = ϵ*_0_, *and the kernel of the linear transformation* 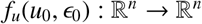 *is spanned by the nonzero vector* 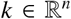. *If* 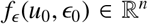 *and* 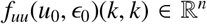 *are both nonzero and both not in the range of f_u_*(*u*_0_, *ϵ*_0_), *then there is a saddle-node bifurcation at u = u*_0_. *Here we define f_uu_*(*u*_0_, *ϵ*_0_)(*k, k*) *as the second Fréchet derivative of f evaluated at* (*u*_0_, *ϵ*_0_) *in the directions given by k and k*.

## 2 Results

### 2.1 Bistability in a biterminal Futile Signaling Cycle

Using the general technique established in the Methods section, we will continue by considering a non-uniterminal example. For this example we will be considering a key reaction network found predominantly in eukaryotic signaling systems, namely a futile signaling cycle, that exhibits bistability when featuring a two-state kinase, as presented in [23]. This reaction network is represented in Figure 4. From Figure 4b, one can see that the linkage class 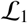 has two terminal strong linkage classes (due to reactions *r*_3_ and *r*_6_). Thus, we have a biterminal linkage class and we cannot apply the theory presented in [18]. Letting

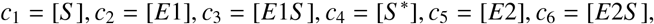

and performing the steps outlined in the Methods section, we obtain the following independent ODEs

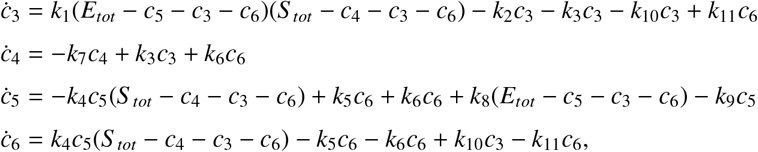

with conservation laws

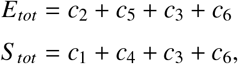

and fix particular kinetic rate constants to expressions (as described in the Methods section) in order to ensure a steady state in the ODE system

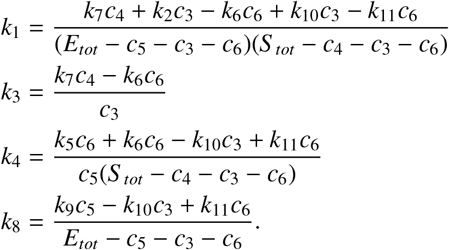

**Fig. 4:**
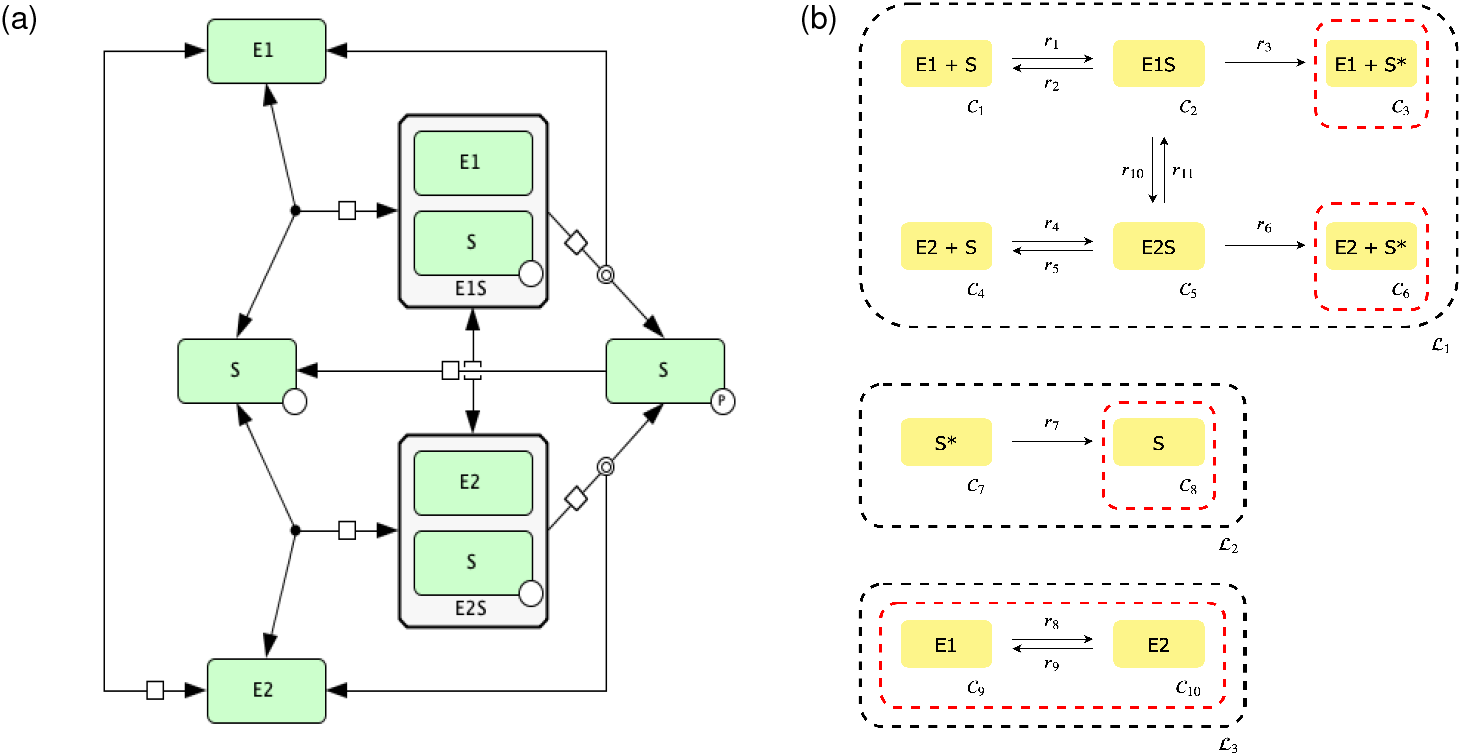
Futile signaling cycle. (a) diagram of reaction network and (b) its C-graph representation.

After performing the optimization routine, we obtain the following values from our decision vector

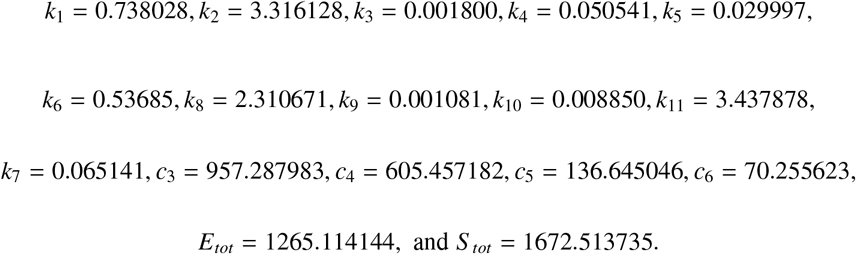

Using the values for the species’ concentrations and kinetic rate constants found, we can now begin to construct the dose-response curve. This is done by using several different concentrations of species *S*\* for the initial value and simulating the ODE system until it has reached a steady state. For this particular example, we obtain the dose-response diagram presented in Figure 5.

**Fig. 5:**
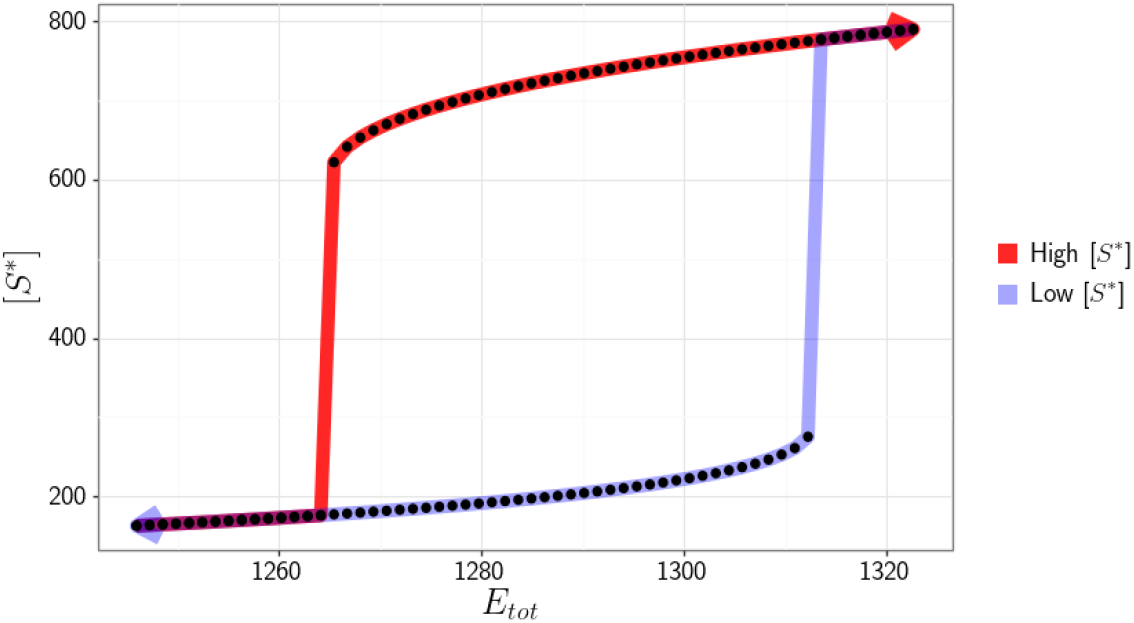
Dose-response diagram exhibiting switch-like behavior produced by a futile signaling cycle. Dots represent the equilibrium that the system converges to for individual simulations. The red and light blue paths correspond to high and low initial concentrations of [*S*\*], respectively.

### 2.2 Bistability in a biterminal Prion/Double Phosphorylation motif

Next we consider the hypothetical mechanism for prion-like conformation conversion between two states of a protein described in Figure 6. Kinase can be in two conformations E1 and E2. Conversion between E1/E2 proceeds through a prion-like mechanism, that is catalyzed by the enzyme E in the corresponding conformation. Only one conformation of kinase, E2, is active and phosphorylates substrate S in two-steps. From inspection, one can see that 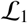 is a linkage class with two terminal strong linkage classes (due to reactions *r*_5_ and *r*_6_). Thus, we have a biterminal linkage class and we cannot apply the theory presented in [18]. Letting

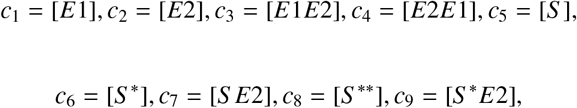

and performing the steps outlined in the Methods section, we obtain the following independent ODEs

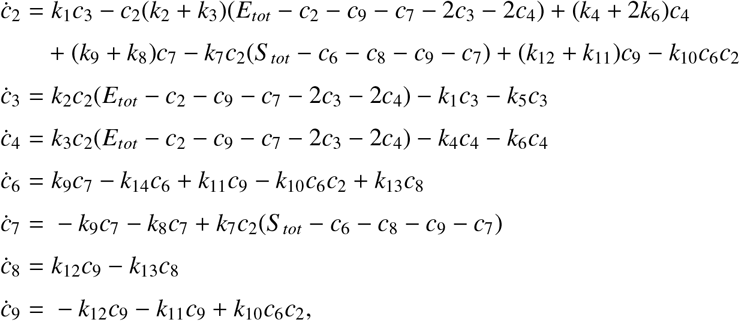

with conservation laws

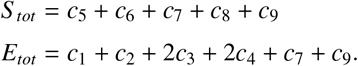

**Fig. 6:**
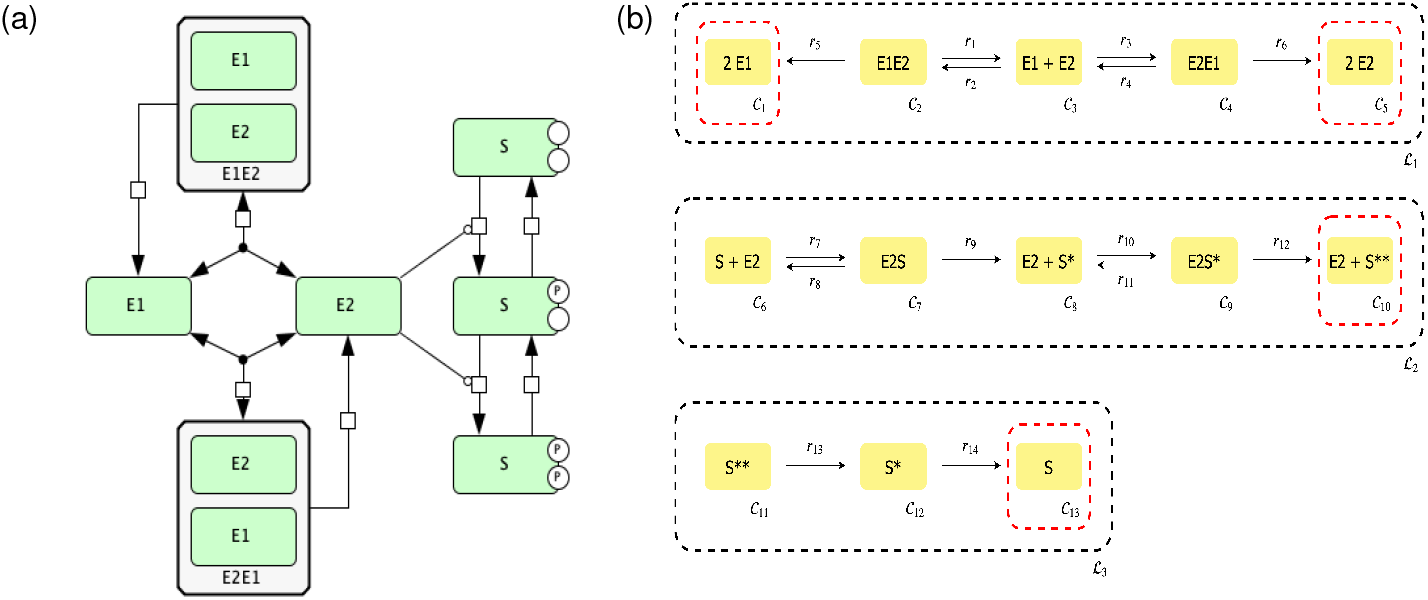
Prion/Double Phosphorylation motif with prion-like conformation conversion between the two states of the same protein. One of the conformations is assumed to be an active kinase. (a) diagram of reaction network and (b) its C-graph representation.

An alternative way to ensure the steady state in the optimization problem is to directly enforce the time derivatives of the concentrations to be zero. This approach can be helpful in cases where the Jacobian is ill-conditioned. Here, for demonstration purposes we choose not to fix the kinetic rate constants and instead use the aforementioned robust, but more computationally intensive approach. Performing the optimization routine, we obtain the following values from our decision vector

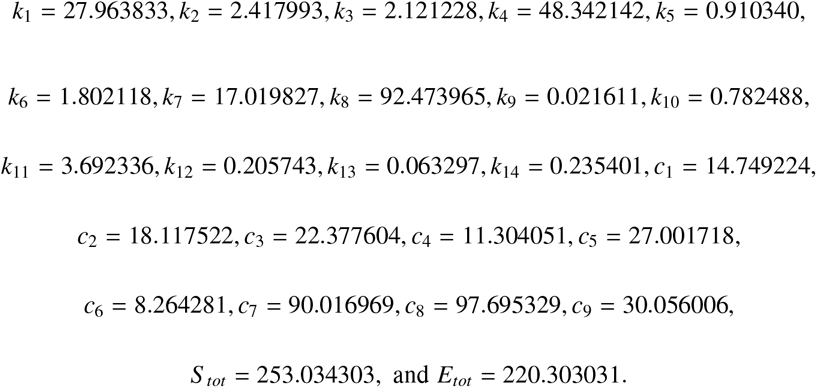

Using the values for the species’ concentrations and kinetic rate constants found, we can now directly compute the bifurcation diagram with continuation methods or alternatively obtain the dose-response curve by direct simulation. This is done by using several different concentrations of species *S*\*\* for the initial value and simulating the ODE system until it has reached a steady state. For this particular example, we obtain the dose-response diagram presented in Figure 7.

**Fig. 7:**
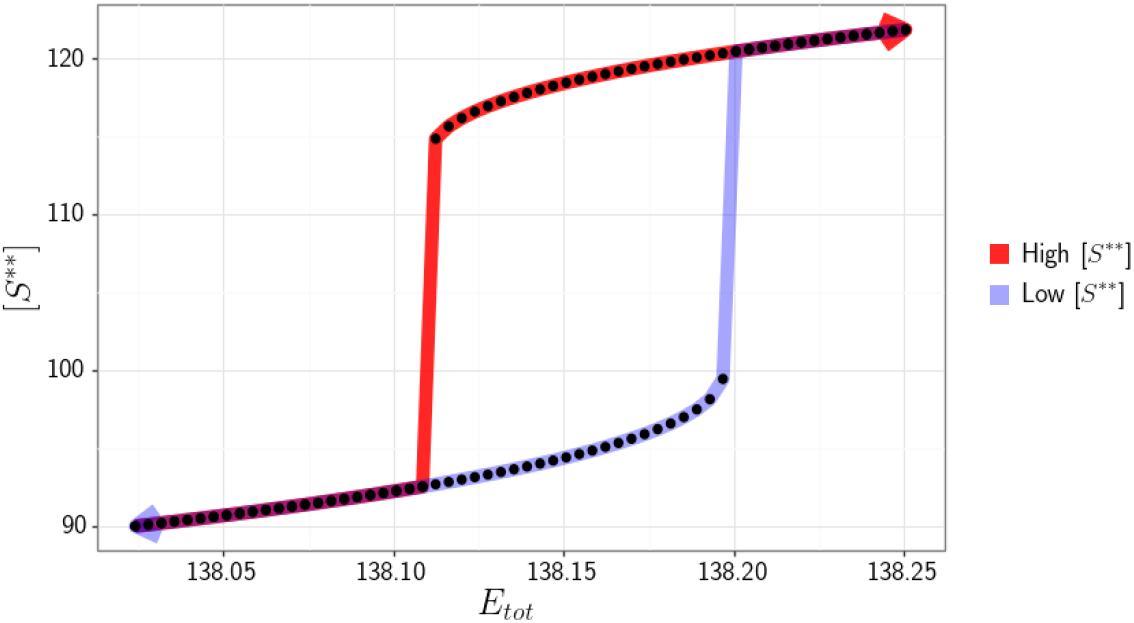
Dose-response diagram exhibiting switch-like behavior produced by the Prion/Double Phosphorylation model.

## 3 Conclusions

We have developed a new general technique to detect bistability in any mass conserving reaction network. The general technique established builds on an existing approach that only allows for bistability detection in reaction networks that are uniterminal. This is achieved by first constructing an objective function using CRNT that is not dependent on the number of terminal strong linkage classes in each linkage class. Then, we formulate an optimization approach that searches for a saddle-node. Once a saddle-node is detected, the method performs either numerical continuation or direct simulation to identify if the particular set of parameters that produced the saddle-node also produce a saddle-node bifurcation. Lastly, if a saddle-node bifurcation is found, numerical continuation or direct simulation will elucidate whether or not there is another saddle-node bifurcation that jointly form switch-like behavior. Various examples provided verify the general technique and its ability to identify bistability. This technique in its entirety is available in an updated version of CRNT4SBML, a Python package that analyzes SBML files (the systems biology community standard for representing reaction networks) and then utilizes mathematical theories to help detect the existence of bistability in cell signaling pathways [24].

Although the general technique is a useful method for the detection of bistability, there are certain difficulties it may encounter.The technique can become computationally intensive for large reaction networks with very high dimensional search spaces to be explored. To overcome this, we included an option that allows the user to perform the optimization routine in parallel. Additionally, even if a large portion of the objective function’s domain is explored, we cannot preclude bistability if a zero is not found. We address this uncertainty by computing the probability that the minimum objective function value achieved is equal to the true global minimum. In the future, we would like to couple the general technique with other methods that utilize the Gröbner basis for the ODE system. This coupling could produce further insight into the conditions of bistability for particular parameters produced by the optimization.

## 4 Methods

The introduced approach is more general than those methods developed in [18] as it does not require the network to be uniterminal. Without loss of generality and for the sake of simplicity, we will be using the well-known Edelstein network, which is uniterminal. Originally, the Edelstein network was introduced in [25] and we choose to use the reduced form presented in Lecture 3 of [26]. The Edelstein network is shown in Figure 8. In the following subsections we develop our framework using aspects of CRNT. Here we have **k** = (*k*_1_, *k*_2_, *k*_3_, *k*_4_, *k*_5_, *k*_6_)^*T*^ and **c** = (*c*_1_, *c*_2_, *c*_3_)^*T*^ with *c*_1_ = [*A*], *c*_2_ = [*B*], *c*_3_ = [*C*].

**Fig. 8:**
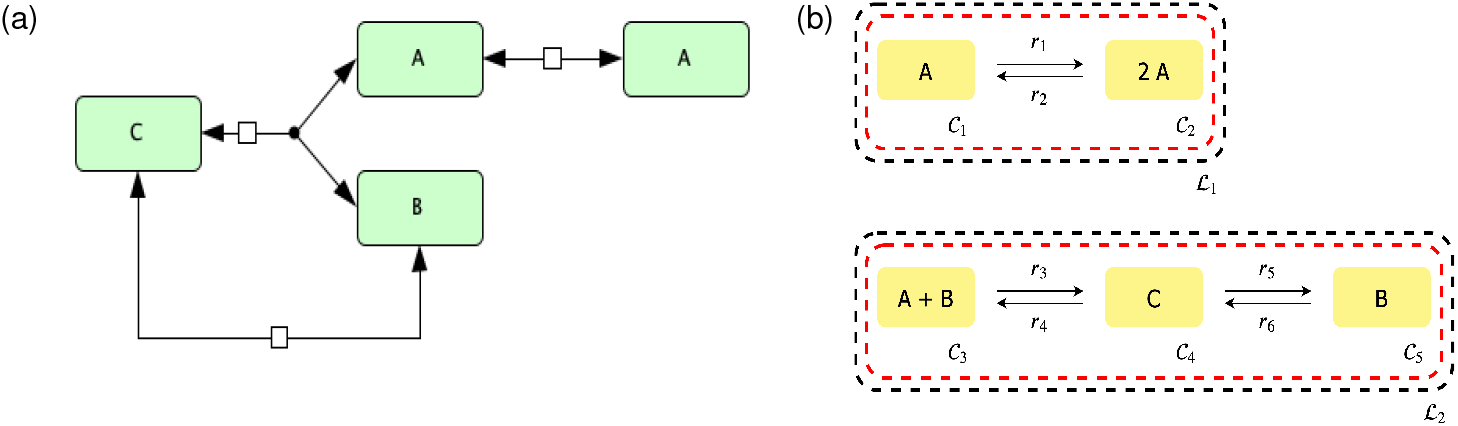
Edelstein chemical reaction network [25]. (a) diagram of reaction network and (b) its C-graph representation.

### 4.1 Step 1: *Constructing the full ODE system*

The first step of the approach is to construct the ODEs describing species’ concentration dynamics for the given reaction network. Assuming mass action kinetics, we obtain the following representation for our autonomous ODEs:

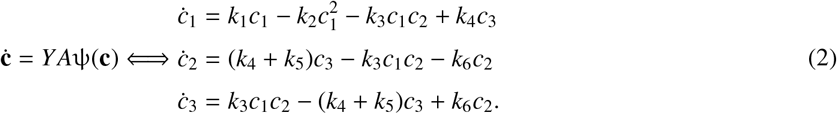

The molecularity matrix *Y* is a *N × M* matrix where *Y_ij_* corresponds to the molecularity of species *i* in complex *j*. For our example the complexes are *C*_1_ = *A*, *C*_2_ = 2*A, C*_3_ = *A* + *B, C*_4_ = *C*, and *C*_5_ = *B*, which provide the molecularity matrix:

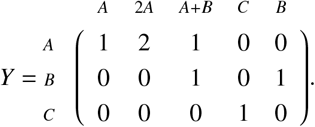

*A* is the *M × M* kinetic constant matrix, where the diagonal elements of *A* contain the negative of the sum of the kinetic rate constants corresponding to the reactions going out of *C_i_*, while the off-diagonal elements contain the kinetic rate constants of the reactions going from *C_j_* to *C_i_*. Here *i* and *j* correspond to the *i*th row and *j*th column of A, respectively

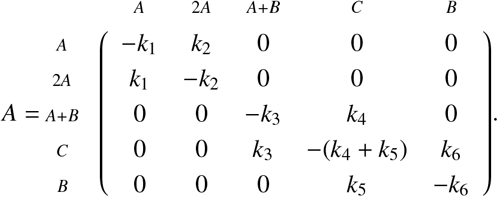

The vector ψ(**c**) = (ψ_1_,…,ψ_*M*_)^*T*^ defines the mass action monomials (a product of species’ concentrations) associated with each complex

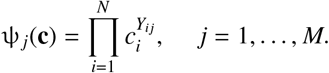

For our example we obtain:

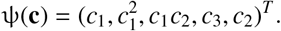

### 4.2 Step 2: *Computing the mass conservation laws*

The next step is to compute the mass conservation laws that govern the network. To do this, we first construct the species stoichiometric matrix *S* with dimension *N × R*. Its entries can be constructed from the *Y* matrix by noticing that every reaction from *C_i_* to *C_j_* has an associated vector *Y_j_ – Y_i_* where *Y_j_* and *Y_i_* are the *j*th and *i*th columns of Y, respectively. In our example, this produces the stoichiometric matrix *S*:

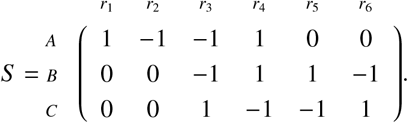

We also denote the rank of *S* as *s*, for notational convenience. For our example, *s* = 2.

Note, that the number of mass conservation relations is *λ = N – s*. Matrix *B* (of dimension *N × λ*) that defines such relations is defined as the nullspace of *S^T^*, such that *S^T^B* = 0. It should be also noted that the matrix spanning the nullspace of *S^T^* is not uniquely determined, and we choose *B* such that all of its entries are nonnegative. This choice of *B* is always possible provided that each conservation law represents the conservation of a chemical or moiety [18].

For most reaction networks, constructing a *B* matrix with nonnegative entries can be done by using linear programming. In particular, an optimization problem is formed by first considering the intersection of *Null*(*S^T^*) and the nonnegative vectors of 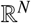, which is an infinite cone. In order to obtain a finite set of this intersection, one can consider an intersection of the infinite cone with the set of vectors that have the sum of their coordinates equal to one. The solution to this intersection is a finite convex polytope. Using this form one can search for the vertices of this convex polytope beginning from random directions via the optimization problem (3) solved by the Simplex optimization method [27]. Where **w**^*T*^ is the vector of random search directions and 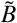 is the initial *B* matrix with at least one negative entry. The endpoints (i.e. the minimized **x** from (3)) of these vertices then form a linear basis element of the original nullspace consisting of only a nonnegative vector. Thus, by taking the dot product, 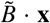, we can obtain a vector with nonegative entries, with values between zero and one. To obtain integer entries, one can then divide the vector by the smallest nonzero entry of the vector. By repeating this process multiple times and obtaining λ unique basis elements, one can then form a *B* matrix with nonnegative entries, as outlined in Algorithm 1. In addition to this approach, there exists alternative ways to obtain a *B* matrix with nonnegative entries, see [28, 29, 30].

Minimize ***w^T^* x** subject to:

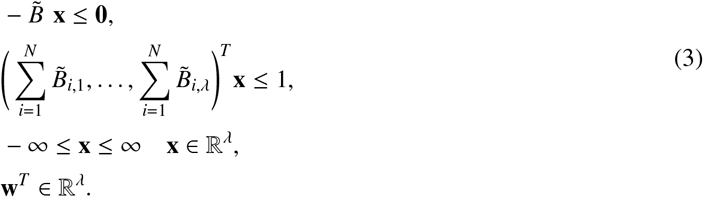

Set 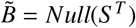.

Set the number of random search directions to *niter* = (*N* + 1)(*λ* + 1)10.

**Algorithm 1:**
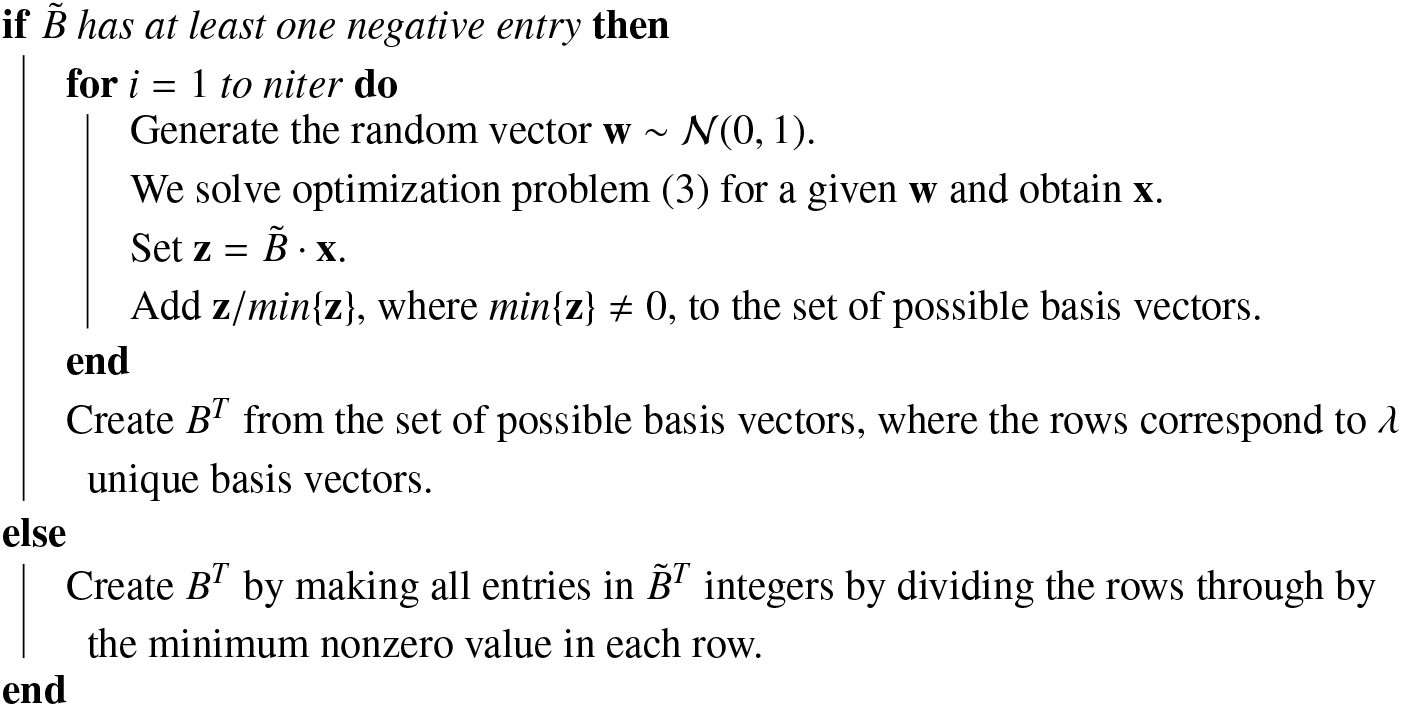
Pseudocode for obtaining a *B* matrix with nonnegative entries.

Using *S^T^* and the procedure outlined in Algorithm 1, we obtain the *B* matrix below:

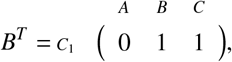

where *C*_1_ again stands for the first conservation law. To obtain an explicit statement for the conservation laws, we simply take *B^T^***c**, giving the following conservation law:

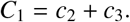

### 4.3 Step 3: *Determining the independent ODE system*

For any given reaction network, the number of independent ODEs describing the system’s dynamics is equal to the rank(*S*) = *s ≤ R* [9]. Since we will ultimately consider the Jacobian of the ODE system with respect to the concentrations **c**, it is necessary to remove those ODEs that can be represented as linear combinations of other ODEs in the system. We will let the independent system of ODEs be represented as follows:

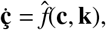

where 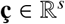 represents those species’ concentrations that form an independent ODE system. To obtain the independent system, we will first see if *s = N*, if this is true, this implies that there are no conservation laws. This scenario is out of the scope of this manuscript. If *s < N* then we have conservation laws and we must continue by finding the independent system. It is at this point where knowledge of the response in the bifurcation analysis is used to determine which ODEs should remain in the independent ODE system.

For our particular example, we take *C*_1_ (the sum of *c*_2_ and c_3_) as the input signal, and *c*_1_ as the output or readout of the system’s response. Since *c*_1_ (the concentration of species A) is taken as our system’s response, it is necessary for *ċ*_1_ to be included in 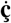. Using this fact and conservation law *C*_1_, we can see that *c*_2_ has a dependence on *c*_3_ of the form *c*_2_ = *C*_1_ − *c*_3_. For this simple example, this information is sufficient enough to eliminate *ċ*_2_ from **ċ**. Indeed if we consider the full ODE system (2), we see that *ċ*_2_ = −*ċ*_3_. Thus, for our particular example we have 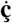 given by:

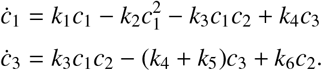

For larger systems the above methodology may not be straightforward. In practice, a set procedure to find this independent ODE system can be conducted. The pseudocode for this procedure is provided in Algorithm 2.

Let *y* be the desired response species.

Let *PDS* be a list of length *λ* initialized with empty lists as its elements.

**Algorithm 2:**
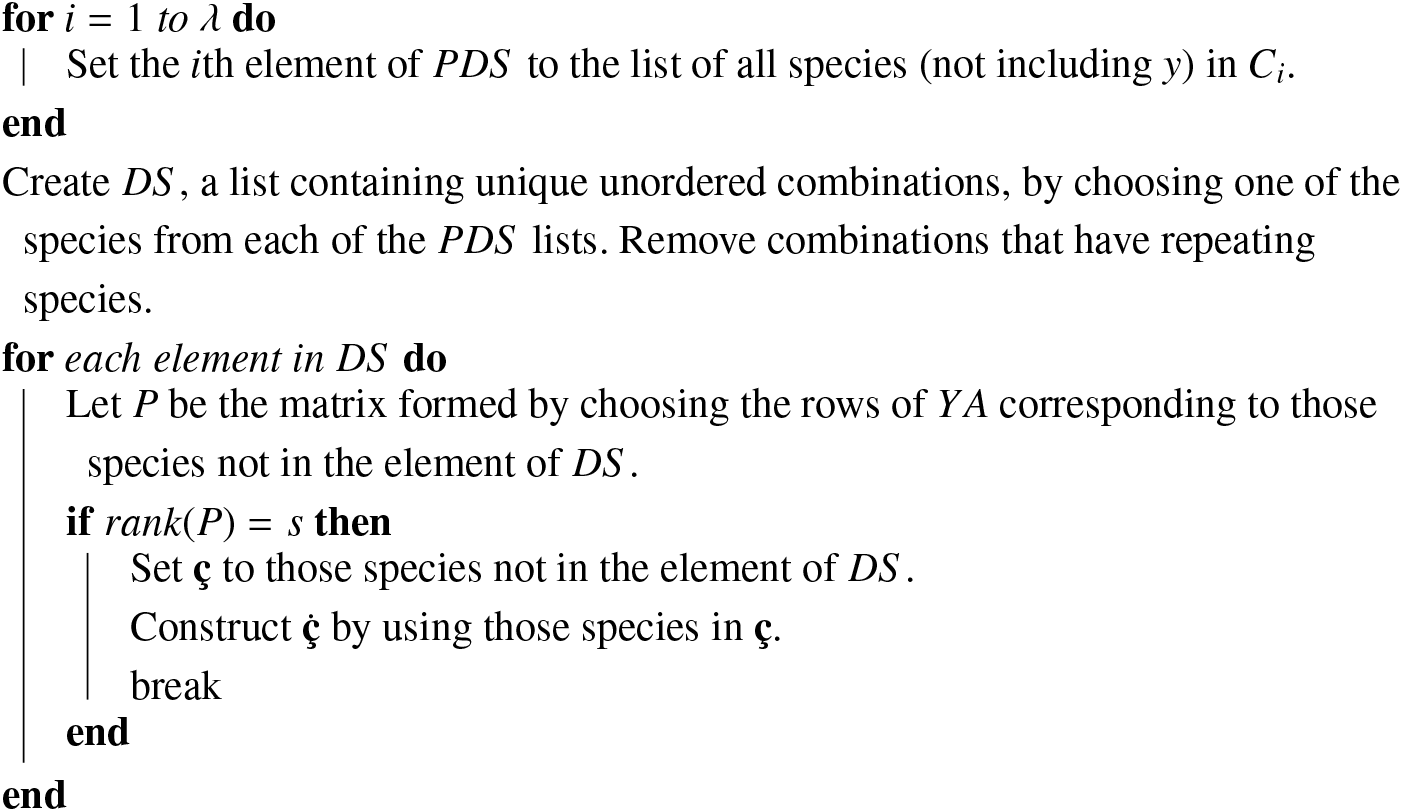
Pseudocode for finding a independent ODE system. Where *PDS* denotes the lists of possible dependent species for each conservation law and *DS* contains all lists that represent our dependent species.

Referring to Definition 4, we see that *u* and *ϵ*_0_ correspond to **c** and the input signal in our example, respectively, with *u* = (*c*_1_, *c*_3_)^*T*^ and *ϵ*_0_ = *C*_1_ From this definition we see that some modifications to 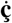 must be made, in particular, we need to include the conservation law *C*_1_. To eliminate all the dependent species (DS) we express them in terms of **ç** and the corresponding conservation law. In our example, this is done by replacing *c*_2_ with *C*_1_ − *c*_3_.

For our example, we then have 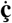 given as follows:

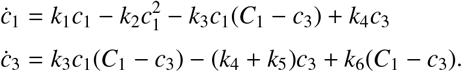

### 4.4 Step 4: *Preserving the steady state*

Now that we have our ODE system in terms of our input signal (or bifurcation parameter), we can now continue to construct those variables that are necessary to check the requirements of Definition 4. To do this, the first item we consider is the statement that *f*(*u*_0_, *ϵ*_0_) = 0, this denotes that the ODE system is at a steady state. Thus, we must ensure that 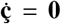. In practice one can formulate this as an optimization problem. Although this formulation is robust, the trade-off includes increased computational time.

To avoid this issue, we suggest an alternative formulation of the optimization problem. We analytically solve for specific kinetic constants, 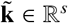, to enforce a steady state. By solving for these expressions analytically, the domain of feasible points is reduced, providing a simpler optimization problem. This can be performed systematically because we are using mass action kinetics to form our ODEs. Furthermore, we are always guaranteed a unique solution in terms of a particular choice of kinetic constants. To see this, consider the stoichiometic matrix *S*, which has columns corresponding to the kinetic constants. As stated in step 3, rank(*S*) = *s ≤ R*, for any given reaction network. By definition, s corresponds to the number of linearly independent columns of *S* and since the columns of *S* correspond to the kinetic constants, we are provided with a set of kinetic constants that form a linearly independent system of equations. In addition, we are also guaranteed to have enough reactions to form this system from our independent ODE system because *s ≤ R*.

In practice, it is quite simple to determine the expressions for 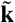, that should be chosen to create a linear system. One particular way to choose these kinetic rate constants is to put *S* into row reduced echelon form (RREF) and choose those columns that contain pivots. In our example:

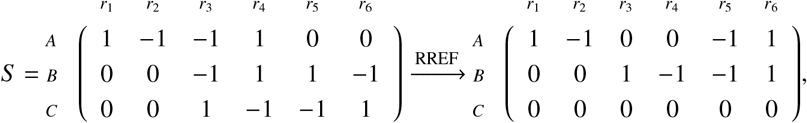

the pivots are in the columns 1 and 3, resulting in 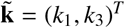.

Given we are looking for a steady state, we must consider the following linear system:

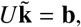

Here 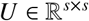 corresponds to the coefficient matrix produced from 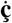 for a particular choice of 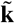. Vector 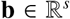 are those terms of 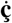 that do not contain kinetic constants from 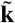. For our example we have the following system for our steady state:

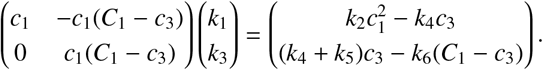

Solving for *k*_1_ and *k*_3_ provides necessary conditions for a steady state

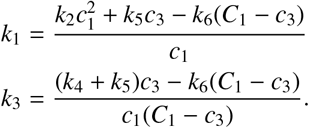

Note that for some systems, this may force a kinetic rate constant equal to zero. However, other choices of 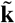 can yield nonzero values. To account for these cases, if zero values are found, we exhaustively search through all distinct combinations of 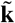 to search for one that yields no zero entries. If no combination provides nonzero values this means that no steady state exists when **k** > 0 and **c** > 0. In such a scenario we suggest checking the Deficiency Zero and One theorems [26]. To obtain a network that can assume a steady state with **k** > 0 and **c** > 0, one can consider removing the reactions that have kinetic constants equal to zero in the aforementioned computation.

### 4.5 Step 5: *Ensuring a zero eigenvalue*

From step 4, we have necessary conditions for a steady state. Thus, the next item we must satisfy is that the Jacobian with respect to **ç**, *J_ç_*, must have a zero eigenvalue with a multiplicity of one and all other eigenvalues have nonzero real parts as stated in Definition 4. Given that we would like to formulate this as an optimization problem, it is easy to see that satisfying the criteria that an eigenvalue of zero must have a multiplicity of one and all other eigenvalues be nonzero, will create an expensive optimization problem because we would have to calculate the eigenvalues each time the objective function is evaluated. For this reason, we will simply search for a zero eigenvalue and then check afterwards if the eigenvalue criteria are satisfied.

To find a zero eigenvalue, consider the eigenvalue problem with eigenvalue 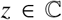 and corresponding eigenvector 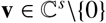:

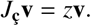

The characteristic polynomial can then be formed by taking the determinant and setting it equal to zero:

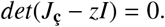

However, since *z* = 0, we have the following problem that reassures us that at least one eigenvalue is equal to zero:

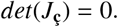

It is this criteria that we will use to formulate our optimization problem.

### 4.6 Step 6: *Searching for a Saddle-node using optimization*

Using the definitions from the previous steps we can create the optimization problem in two alternative ways. In both formulations, the optimization problems search for species’ concentrations **c** and kinetic rate constants **k** that provide 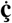 with at least one zero eigenvalue. One that implicitly enforces a steady state by solving for kinetic constants (as in Step 4):

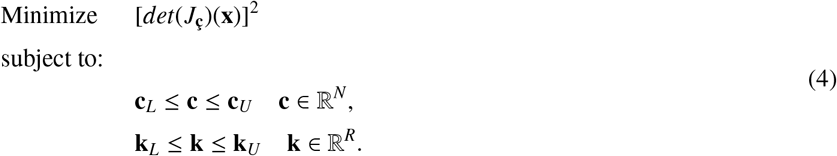

In this version of the optimization problem **c**_*L*_ and **c**_*U*_ are the lower and upper bounds for the species’ concentrations, respectively and similarly, **k**_*L*_ and **k**_*U*_ are the lower and upper bounds for the kinetic rate constants, respectively. The decision vector 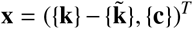, where 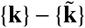 corresponds to the set of kinetic rate constants in **k** that are not in 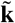 and {**c**} is the set of species’ concentrations. The kinetic rate constants 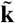 are then found by using the solutions found from solving the linear system in step 4, which reassure a steady state occurs, and the conservation constants *C*_1_,…, *C_λ_* are found using the conservation laws in step 2.

Another formulation enforces a steady state by explicitly requiring that the derivatives of the concentrations be zero through an additional term in the objective function:

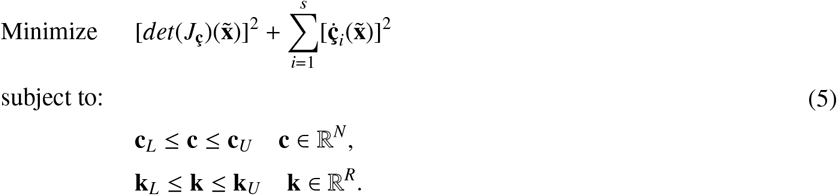

In this alternative version we instead have the decision vector 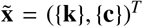. This results in a more complex objective function, which is due to the addition of more nonlinear terms in the objective function and a higher dimensional decision vector. However, this approach is more robust as it samples from a broader space of solutions.

Due to their nature, like the optimization problem in [18], (4) and (5) are non-convex and multi-modal. For this reason, a global optimization procedure should be used to search for the optimal solution of (4) and (5). For the results obtained throughout the manuscript, we utilize the global optimization algorithm Dual Annealing, starting from different random initial starting points that are bounded by the species’ concentrations and kinetic constant values. Dual Annealing is an optimization algorithm that combines Simulated Annealing [31] and additionally applies a local search on accepted locations [32]. In particular, we utilize the Nelder-Mead simplex algorithm for all local searches. Given that we are looking specifically for a zero eigenvalue, the objective function of the optimization should obtain a minimum of zero. If a zero is found one should additionally provide a check for i) a saddle-node using Definition 4 and discard points that produce more than one eigenvalue that is zero (which might indicate a codim 2 bifurcation such as the Bogdanov-Takens bifurcation [33]), and ii) a saddle-node bifurcation point using Theorem 1. This allows one to remove unnecessary runs of numerical continuation. Although explicitly checking the criteria for a saddle-node bifurcation can reduce the runtime of the approach, it should be noted that it is sufficient to conduct numerical continuation.

#### 4.6.1 Bayesian Stopping rule

If the optimization satisfies the condition for a saddle-node, one can state that a saddle-node exists. However, the inverse is not true: if we can’t find conditions that result in a zero eigenvalue, this doesn’t guarantee that a saddle-node does not exist. This is a consequence of using stochastic optimization instead of deterministic methods (based for example on interval analysis), which will provide this guarantee [34]. The issue with methods such as these is that they become computationally intractable for mass action reaction systems with more than two or three parameters, whereas stochastic optimization provides in general a good overall efficiency [35]. To address this uncertainty we compute the probability that the achieved minimum value is equal to the true global minimum. The probability is calculated using a slightly modified version of the unified Bayesian stopping rule in [36] and Theorem 4.1 of [37], where the rule was first established.

For *n* starting optimization points, let *f_k_* be the achieved local minimum for the *k*-th decision vector, *f*\* the true global minimum and 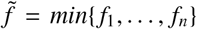. Our objective is to estimate the probability that 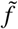 is *f*\*. Let *α_k_* and *α*\* denote the probabilities that a single run of the optimization routine has converged to *f_k_* and *f*\*, respectively. Assuming that *α* ≥ α_k_* for all local minimum values *f_k_* we may then estimate the lower bound of the probability that 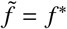 is as follows:

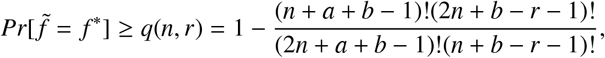

Here *a* and *b* are the parameters of the Beta distribution *β*(*a, b*), where we use *a* = 1 and *b* = 5 as suggested in [36]. The term *q*(*n, r*) is the confidence level, where *r* is the number of *f_k_* for *k* = 1, …, *n* that are in the neighborhood of 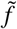.

We say that *f_k_* is in the neighborhood of 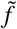 if the relative difference of *f_k_* and 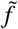 is less than 1%:

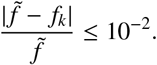

Given the formulation of the optimization, the lowest possible minimal value is zero. Thus, if 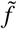 reaches a numerical zero value then we set *q*(*n, r*) = 1.0, skipping the computation of *q*(*n, r*). Conventionally *q*(*n, r*) ≥ 0.95 is considered an acceptable confidence level to make the conclusion that 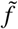 is the global minimum of the objective function.

#### 4.6.2 Demonstrating the Bayesian Stopping Rule

Now that we have established an approach to estimate the probability that the minimum objective function value is the global minimum, we will demonstrate this by using an example. Consider the reaction network provided in Figure 9. By the Deficiency Zero Theorem of [9] it is known that the network cannot be bistable and there is only one equilibrium.

**Fig. 9:**
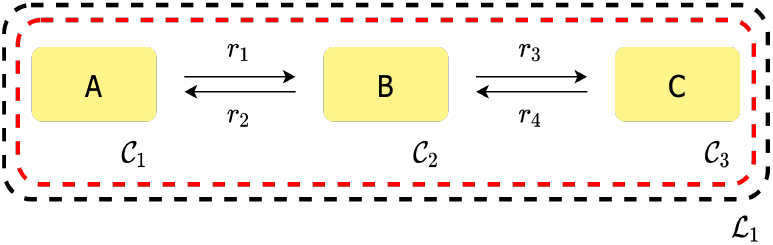
C-graph of a simple non-bistable network.

Because of the simplicity of the reaction network we can compute the exact value of the global minimum (e.g. using Mathematica’s MinValue [38]). Letting our reaction coefficients and species’ concentrations be bounded between 0.01 and 100, the lowest possible value is *f*\* = 9.0*e* − 8. Performing 100 iterations of the optimization routine, we obtain 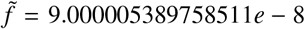 with a confidence level of *q*(*n, r*) = 0.9999999995824618. Since *q*(*n, r*) is greater than 0.95, we would conclude that the achieved value of 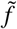 is the global minimum. For this reason, we can reject the possibility that a zero value of the objective function can be achieved.

### 4.7 Step 7: *Numerical Continuation and Direct Simulation*

Once the optimization problem is solved, and the saddle-node condition checked, we have an independent ODE system with a steady state and exactly one zero eigenvalue. To check if these particular parameters produce a saddle-node bifurcation point, we either utilize the technique of numerical continuation or directly simulate the ODEs. When performing numerical continuation, we use the values provided by the optimization routine and then leverage the established tool AUTO 2000 [39]. It is made accessible through the libroadrunner python library and its extension rrplugins [40]. To demonstrate numerical continuation, we use the following values provided by the optimization routine, which provide Figure 10:

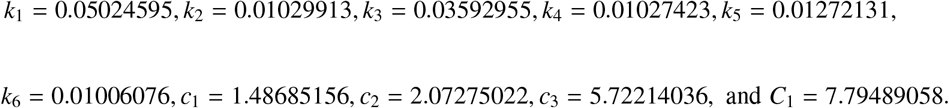

**Fig. 10:**
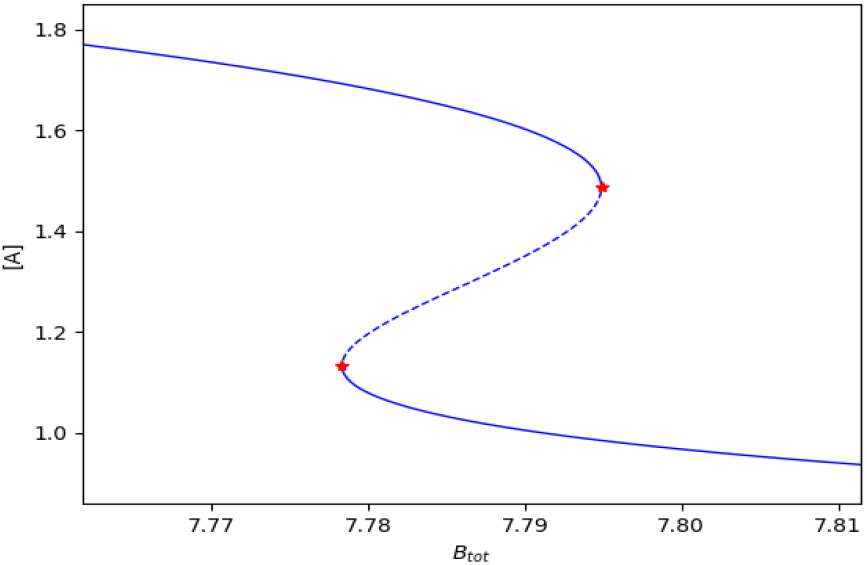
Bifurcation diagram of the Edelstein network example. The bifurcation diagram was created using the software AUTO 2000, which utilizes numerical continuation. Red markers represent the found saddle-node bifurcation points, the stable branches are solid blue lines, and the unstable branch is a dashed blue line.

It should be noted that it is possible that the optimization routine finds kinetic rate constants that force the Jacobian of the system to be ill-conditioned or even singular, even if species concentrations are varied. If this particular scenario occurs, numerical continuation will not be able to continue as it relies on a well-conditioned Jacobian. To overcome this type of situation we offer the option of direct simulation, which varies the user defined signal and initial conditions values for the ODE system and then integrates the ODEs until a steady state occurs. The steady state is obtained when the species’ concentrations do not change between the ODE integration steps, within a predefined tolerance (here 1e-06). Given the direct simulation method is numerically integrating the system of ODEs, this method will often take longer than the numerical continuation routine. Although this is the case, direct simulation is more robust and may be able to provide a bifurcation diagram when numerical continuation cannot.

Considering high and low concentrations of species *A* (response) for the initial value, we obtain the ODE simulations in Figure 11 by direct simulation, where *B_tot_* = *C*_1_.

**Fig. 11:**
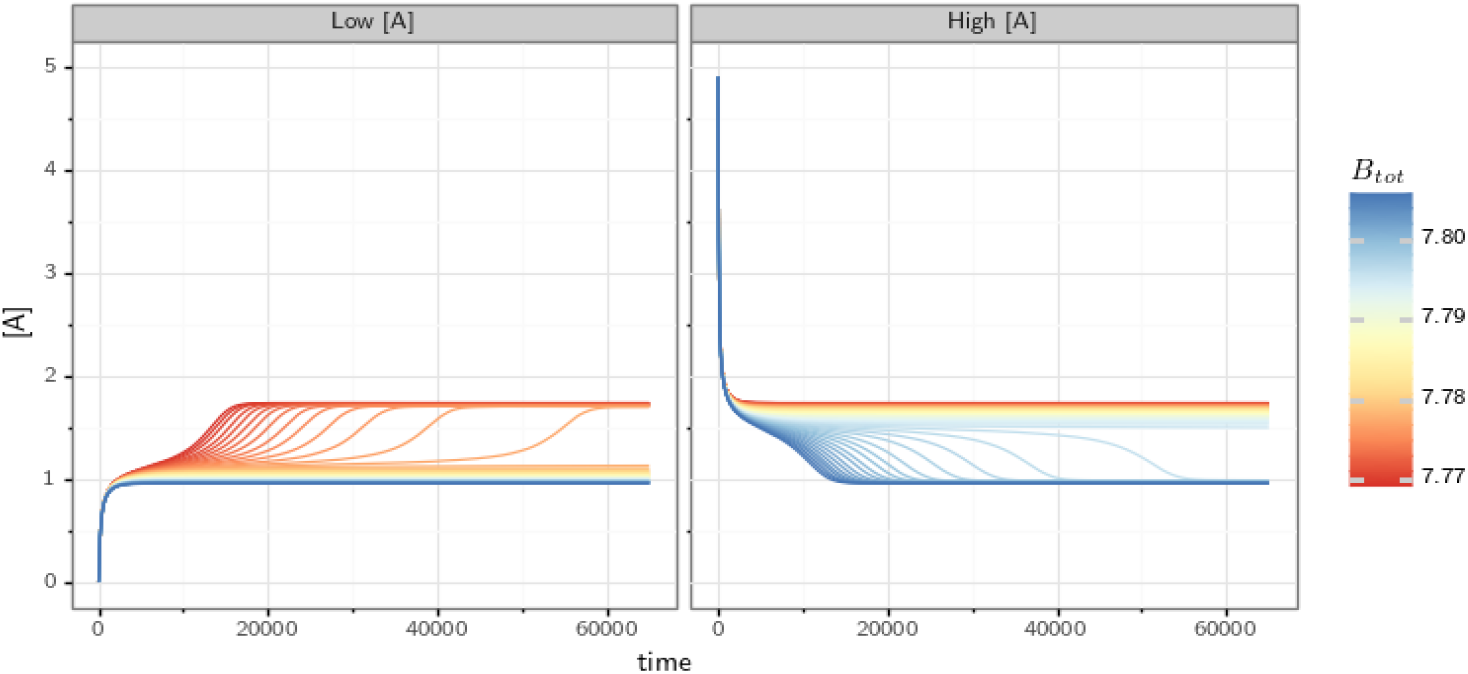
Simulation of the ODE system for the Edelstein network using different starting values of *B_tot_* and [A]. The system converges to two equilibrium states; a “lower” one at [*A*] ≈ 1 and a “higher” one at [*A*] ≈ 1.75. If the system starts with a low concentration of *A* it takes a larger concentration of *B_tot_* to reach the “higher” equilibrium state. Vice versa, if the system starts with a high concentration of *A* then small concentrations of *B_tot_* are sufficient to maintain the “higher” equilibrium state.

If we then consider the left and right plot at time 65,000 as the equilibrium, we can plot the values of *B_tot_* vs the concentration of species *A*. The dose-response diagram obtained by direct simulation depicted in Figure 12 mirrors the bifurcation diagram obtained by numerical continuation in Figure 10, thus cross-validating both approaches.

**Fig. 12:**
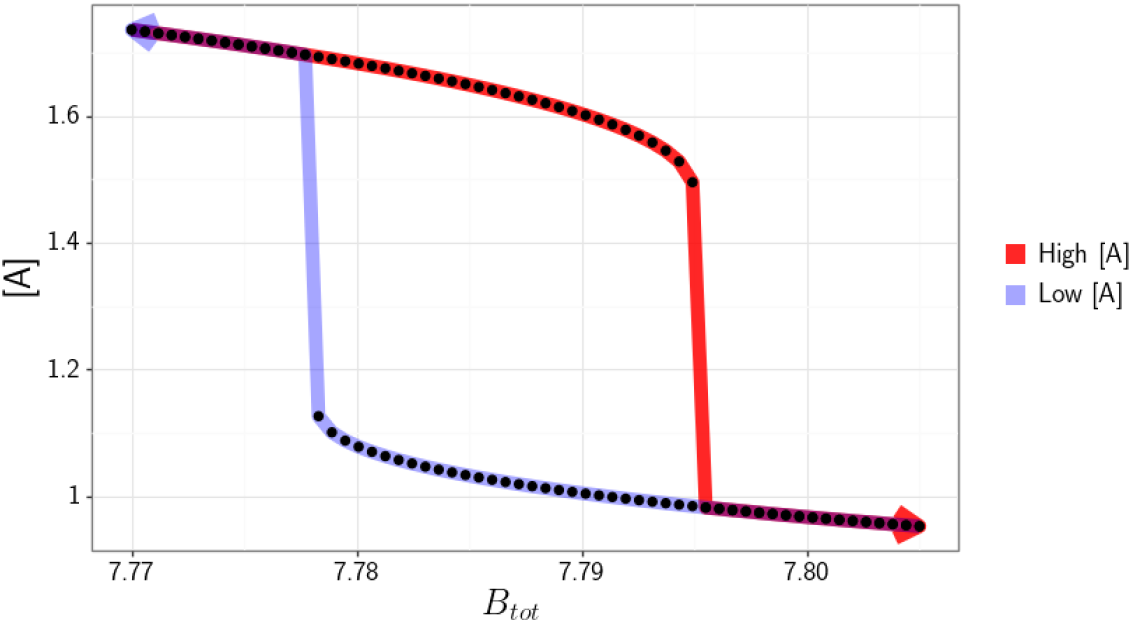
Bifurcation diagram of the example Edelstein network. It is a crosssection of the Figure 11 data at the time point when the system reaches the equilibrium (65,000 seconds for the given parameters). The equilibrium concentration of *A* is plotted for both the “high” and “low” portions of the simulation, highlighted in red and light blue, respectively.

## Author’s contributions

BR and VP developed the methodology. BR developed the code. IOM consulted on the development of the method. All authors wrote, revised and approved the final manuscript.

## Acknowledgements

We would like to thank Matthew E. Monroe for supporting us with computational resources and infrastructure.

